# Reversible edema following electric drilling of macaque craniotomy

**DOI:** 10.1101/2022.03.19.484963

**Authors:** Rober Boshra, Manoj Eradath, Kacie Dougherty, Bichan Wu, Britney M. Morea, Mark Pinsk, Sabine Kastner

**Author notes:** Corresponding authors: Rober Boshra Sabine Kastner.

## Abstract

In-vivo electrophysiology requires direct access to brain tissue, necessitating the development and refinement of surgical procedures and techniques that promote the health and well-being of the animal subjects. Here, we report a series of findings noted on structural magnetic resonance imaging (MRI) scans in monkeys with MRI-compatible implants following small craniotomies that provide access for intracranial electrophysiology. We found distinct brain regions exhibiting hyperintensities in T2-weighted scans that were prominent underneath the sites at which craniotomies had been performed. We interpreted these hyperintensities as edema of the neural tissue and found that they were predominantly present following electric drilling, but not when manual, hand-operated drills were used. Further, the anomalies subsided within 2-3 weeks following surgery. Our report highlights the utility of MRI-compatible implants that promote clinical examination of the animal’s brain, sometimes revealing findings that may go unnoticed when incompatible implants are used. We show replicable differences in outcome when using electric vs. mechanical devices, both ubiquitous in the field. If electric drills are used, our report cautions electrophysiological recordings from tissue directly underneath the craniotomy for the first 2-3 weeks following the procedure due to putative edema.

## Introduction

Research in non-human primates (NHP) is instrumental for probing fundamental questions about the nature and mechanisms of brain function related to perception, action, and cognition. Electrophysiological investigations in NHPs require surgical procedures that enable direct access to the brain in order to target neuronal populations of interest. Techniques to streamline such surgical procedures are continuously developed to optimize the well-being and experimental outcomes of the animal subjects.

Drilling is a core component of the surgical procedures to provide access to neural tissue for electrode placement. Manually-operated tools such as trephines and hand-drills are widely used and require specific technique and skill of the surgeon to ensure that only skull tissue is drilled, and that surgical tools do not compromise the integrity of the dura mater and the underlying brain tissue. In extreme cases, accidental lowering of the drill into tissue, also termed “plunging”, may have severe consequences such as hematomas or irreversible brain lesions ^1–3^. In addition, electric counterparts of the hand-drill are commonly used, as they enable the surgeon to expedite the drilling process. Specifically, in human surgeries, there have been advances in automatic drills that seize upon complete drilling of skull tissue, reducing the possibility of plunging events and clearing bone shavings from the site of drilling ^2,4^. Of note, electric drills that do not feature that safety mechanism are also used in NHP surgeries and require a skilled surgeon to avoid plunging events. Lastly, piezoelectric drills have been introduced as a new class of drills. They utilize ultrasonic vibrations that enable selective removal of hard surfaces (i.e., bone) without harming soft tissue (i.e., the dura or brain tissue). The piezoelectric drill, originally developed for oral surgeries^5,6^, has been adopted for pediatric cranial surgery and has been recommended for NHP craniotomies, as it affords the surgeon a procedure that is less prone to harm brain tissue while being relatively expedient ^7–12^. While different drilling tools have been historically tested in animals and refined in clinical practice for humans, usage in NHP research is quite heterogeneous and may depend on specific laboratory practices or the surgeon’s preferences.

Here, we present a series of magnetic resonance imaging (MRI) scans following 10 craniotomy surgeries with different surgical drilling tools in NHPs implanted with MRI-compatible materials. We found an unexpected anomaly in scans that were collected within two weeks of craniotomies conducted with electric or piezoelectric drills. Specifically, hyperintensities in the vicinity of the craniotomies were noted on T2-weighted scans and were interpreted as edemas. These anomalies were not accompanied by behavioral abnormalities, or any other clinical signs. Our findings have potential implications for intracranial recordings from the tissue directly underlying a craniotomy immediately following a procedure that used an electric drill.

## Methods

### Subjects, inclusion criteria and surgical procedures

Data from eight male macaques (7 *macaca mulatta*, 1 *macaca fascicularis*) were included in the study. All procedures were approved by the Princeton University Animal Care and Use Committee and conformed to the National Institutes of Health guidelines for the humane care and use of laboratory animals. All monkeys were subjects in electrophysiological studies requiring cranial implants and craniotomies to provide access to regions-of-interest in the brain. Each animal had one or two surgeries and was included in the study if they participated in a craniotomy surgery followed by an MRI scanning session within one month post-surgery.

All surgical procedures were performed under general anesthesia with isoflurane (induction 2–5%, maintenance 0.5–2.5%) under strictly aseptic conditions. Vital signs, including heart rate, SpO_2_, respiratory rate, blood pressure, end-tidal carbon dioxide, and body temperature were continuously monitored throughout the procedure. We affixed customized plastic recording chambers to the animals’ skull using ceramic screws (Thomas Recording, Gieseen, Germany in Case 1; or titanium screws, Fine Science Tools; Foster City, CA in cases 9 and 10. Dental acrylic (Yates Motloid Crosslinked Flash Acrylic; Elmhurst, IL), and/or bone cement (Palacos bone cement, PLACE; or DJO Surgical Cobalt HV bone cement, Austin, TX) was used to create a small implant around the chambers and screws ^13^.

In a follow-up surgery, small craniotomies (4.5-8□mm diameter) were drilled inside those chambers. Drilling was conducted in one of four ways: (i) using a set of two mechanical hand-drills (pointed and blunt-tipped; Fine Science Tools) and respective stoppers to penetrate the skull to the dural surface, then complete a cylindrical hole using the blunt-tipped drill (6 surgeries, 13 craniotomies; Table 1); (ii) using an electric drill (RAMPOWER; Ram products inc.) to remove a column of skull tissue entirely leading to the dural surface, removing any thin bone debris using fine forceps (1 surgery, 3 craniotomies; Table 1); (iii) using an electric drill to remove excess cement and to thin out skull tissue prior to using a piezoelectric drill to remove the skull above the dura using lowest drilling setting (endo) and continuous irrigation (1 surgery, 3 craniotomies; Table 1); (iv) strictly using a piezoelectric drill with continuous irrigation for the full craniotomy procedure, starting with an intermediate drilling setting, followed up by the lowest setting once the drill is close to the dural surface (2 surgeries, 6 craniotomies; Table 1). All animal surgeries concluded uneventfully with no visible damage to the dura or plunging events. Following each surgery, animals were monitored closely by facility veterinarians, animal care staff, and researchers. Animals were also administered post-operative medicine, including analgesics and/or antibiotics, following prescription by the veterinarians.

**Table 1:** 10 cases involving 8 animals following craniotomy surgeries with one or more drilling tools, the number of days following the surgery that a scan was conducted, scanner used, and whether T2 hyperintenstities were found. Note that Case 1 had two scans within 1 month post-surgery. M = Monkey, Elec = Electric drill, Piezo = Piezoelectric drill, and Mec = Mechanical drill.

| # | M | Craniotomies | Elec | Piezo | Mec | Scan<br>(days after Sx) | Scanner | Anomalies |
| --- | --- | --- | --- | --- | --- | --- | --- | --- |
| 1 | P | 3 | + | + |  | 7 | Skyra | Yes |
|  |  |  |  |  |  | 14 | Prisma | No |
| 2 | R | 5 |  | + |  | 7 | Prisma | Yes |
| 3 | V | 3 | + |  |  | 6 | Skyra | Yes |
| 4 | Ph | 1 |  | + |  | 22 | Skyra | No |
| 5 | Ph | 3 |  |  | + | 12 | Prisma | No |
| 6 | M | 3 |  |  | + | 5 | Prisma | No |
| 7 | M | 3 |  |  | + | 3 | Skyra | No |
| 8 | F | 1 |  |  | + | 6 | Skyra | No |
| 9 | Mc | 2 |  |  | + | 8 | Skyra | Yes |
| 10 | B | 1 |  |  | + | 4 | Skyra | No |

### Neuroimaging protocols

For scanning sessions, animals were sedated with ketamine (1-10mg/kg i.m.) and xylazine (1-2 mg/kg i.m.), and provided with atropine (0.04 mg/kg i.m.). Sedation was maintained with tiletamine/zolazepam (1-5mg/kg i.m.). The animals were then placed in an MR-compatible stereotaxic frame (1530M; David Kopf Instruments, Tujunga CA). Vital signs were monitored with wireless ECG, pulse, respiration sensors (Siemens AG, Berlin), and a fiber optic temperature probe (FOTS100; Biopac Systems Inc, Goleta CA). Body temperature was maintained with blankets and a warm water re-circulating pump (TP600; Stryker Corp, Kalamazoo MI).

All animals had whole-brain structural MRI data collected either on a Siemens 3 Tesla MAGNETOM Skyra or on a Siemens 3 Tesla MAGNETOM Prisma using a Siemens 11-cm loop coil placed over the head (see Table 1). T2-weighted volumes were acquired with a 3D turbo spin echo with variable flip-angle echo trains (3D T2-SPACE) sequence (voxel size: 0.5□mm, slice orientation: sagittal, slice thickness: 0.5 mm, field of view (FoV): 128 × 128 mm, FoV phase: 79.7%, repetition time (TR): 3390 ms, echo time (TE): 386 ms (Skyra) or 387 (Prisma), base resolution: 256 × 256, and acquisition time (TA): 17 min 41 s (Skyra) or 15:39 (Prisma). T1-weighted structural images were collected using the 3D Magnetization-Prepared Rapid-Acquisition Gradient Echo (MPRAGE) sequence, voxel size: 0.5 mm, slice orientation: sagittal, slice thickness: 0.5□mm, FoV: 128 × 128 mm, FoV phase: 100%, TR: 2700 ms, TE: 3.27 ms (Skyra) or 2.78 ms (Prisma), inversion time (TI): 850 ms (Skyra) or 861 ms (Prisma), base resolution: 256 × 256, and TA: 11 min 31 s.

All animals had scan sessions conducted prior to their craniotomy surgeries for accurate positioning of recording chambers. The scans were inspected by experienced personnel to rule out anomalous findings prior to surgery and served as reference images. Further scans were conducted following the animals’ craniotomy surgeries either for clinical purposes (Case 1, monkey P) or to confirm electrode localizations in brain regions-of-interest.

## Results

### Behavioral signs

Exactly one week following his craniotomy surgery, monkey P was reported for inappetence and reduced social interactions with humans. Precautionary T1- and T2-weighted scans were conducted the following day to ensure integrity of the brain post-surgery. The other animals whose data were included in this report did not show behavioral abnormalities post-surgery.

### Neuroimaging findings and their resolution

Monkey P (Case 1) had undergone three craniotomies in anterior, central, and posterior chambers of the left hemisphere, targeting three regions of interest for electrophysiological recordings from the frontal eye fields, pulvinar, and superior colliculus, respectively. Drilling had been done initially using an electric drill for each craniotomy. Once a thin skull layer was separating the drill from the dural surface, the surgeon switched to a piezoelectric drill set with sufficient irrigation and slow speed (see methods; Table 1, Case 1). The T2-weighted scan taken one week post-surgery showed two white matter hyperintensities located underneath the central and anterior chambers (Fig. 1A, top). The locations of these anomalies were confirmed using stereotaxic coordinates and by reference to anatomical landmarks in comparison with scans obtained pre-operatively to be directly underneath the anterior and central craniotomies. No anomalies were visible underneath the posterior craniotomy. The anomalies were less salient on the T1 scan, but could be detected post-hoc as hypointensities (Fig 1A, bottom). Follow-up scans were acquired one week later and revealed a complete resolution of the anomalies associated with both the anterior and central locations (see Fig 1B). T2 hyperintensities are typically observed in edematous neural tissue along with hypointensive findings on T1-weighted scans^14^ (see Fig 1A, bottom). Although the T1/T2 differential intensity profile would not rule out an acute hemorrhage^15^ or infection^16^, the delineated localized hyperintensities on the T2 scan, the localized hypointensities on the T1 scan, and the fact that the anomalies were directly underneath the craniotomies and confined to their extent led us to interpret the anomalies as edemas of neural tissue caused by cranial drilling. The accidental finding in monkey P motivated a retrospective examination of scans from seven other animals, who underwent craniotomy surgery and had scans acquired within one month post-surgery (see methods and Table 1). We were particularly interested to investigate whether these anomalies were associated with a specific drilling method.

**Figure 1:**
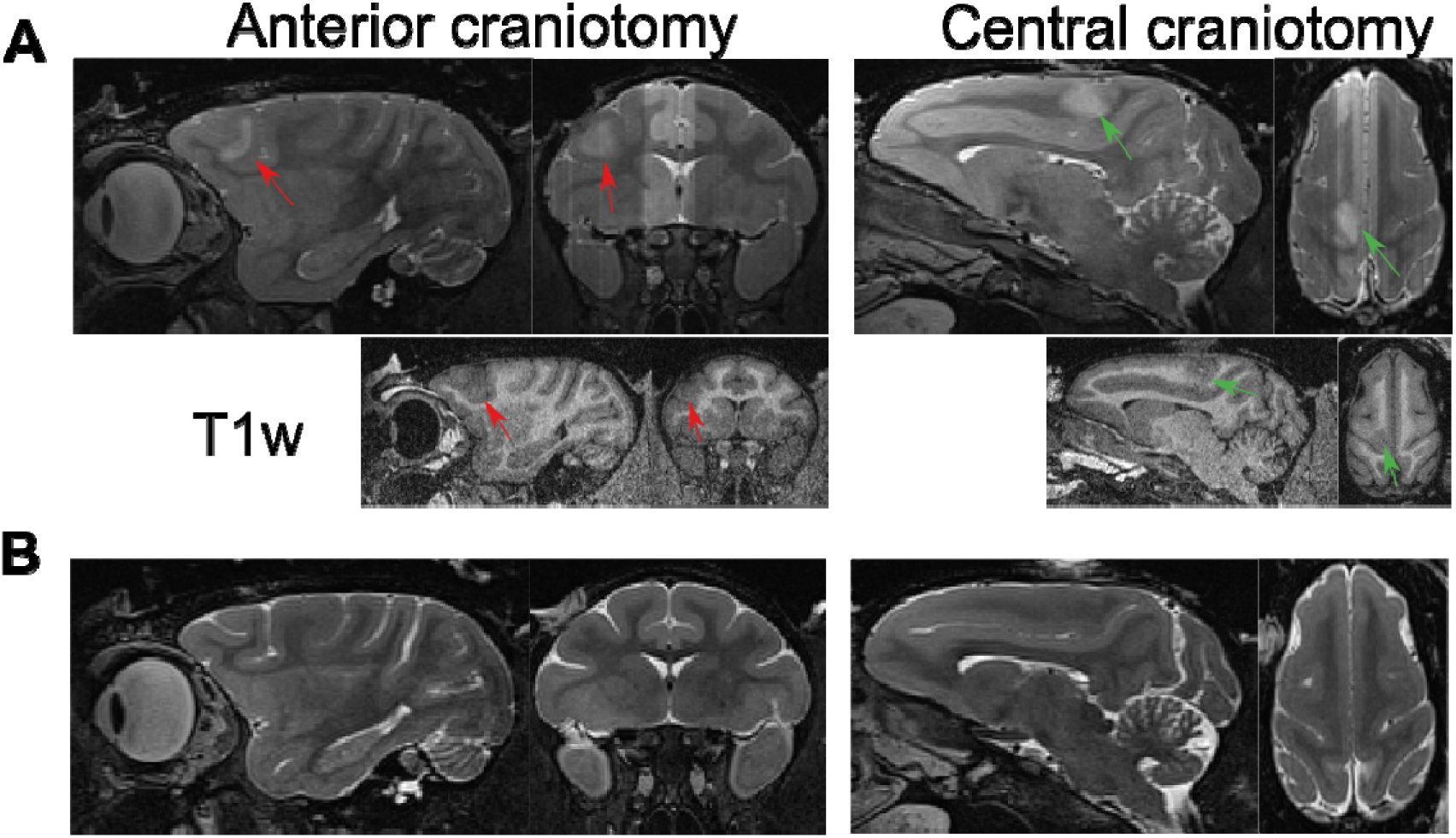
MRI findings in monkey P. White matter T2 hyperintensities found 7 days after surgery in the vicinity of anterior (left; red arrows) and central (right; green arrows) craniotomies in Case 1 (A; top). The same areas show hypointensities on a T1- weighted scan acquired in the same session (A; bottom). These anomalies have resolved on a follow-up scan conducted one week later (B).

Similar T2 anomalies were readily found during our retrospective examination of scans following surgeries utilizing electric and piezoelectric drilling (three surgeries in three animals to drill nine craniotomies; excluding Case 1). Case 2 denotes a surgery that targeted five craniotomies (two in close proximity within an anterior chamber; three in close proximity within a central chamber) in the right hemisphere, which were all performed using piezoelectric drilling. The animal was scanned one week post-surgery. Examination of the scans shows two extensive hyperintensities below the two chambers (Fig 2A). Accurate distinction of whether all craniotomies induced lesioning was not possible, although it is apparent that at least one did not given that an electrode’s shadow did not intersect with the hyperintensity region. The next MR scanning for this animal occurred more than three months later and only included a T1-scan. No trace of characteristic T1 hypointensities under the craniotomies were noted (data not shown). In Case 3, three craniotomies were performed (two in an anterior chamber; one in a central chamber). All drilling in the third case was done exclusively using the electric drill. The T2-weighted scans showed defined hyperintensities (Fig 2B). Similar to Case 2, it was difficult to delineate whether all three craniotomies induced lesions due to the spatial proximity of two of the craniotomies. Case 4 represents the only instance where no anomalous findings on T2-weighted scans were present following craniotomy surgery that used an electric/piezoelectric drill. However, in this case, the animal was not scanned until three weeks following surgery. Based on the profile of the anomalies in our animals and their progression timeline observed in Case 1, it is possible that, if an edema was induced during the surgery, it may have resolved at this point.

**Figure 2:**
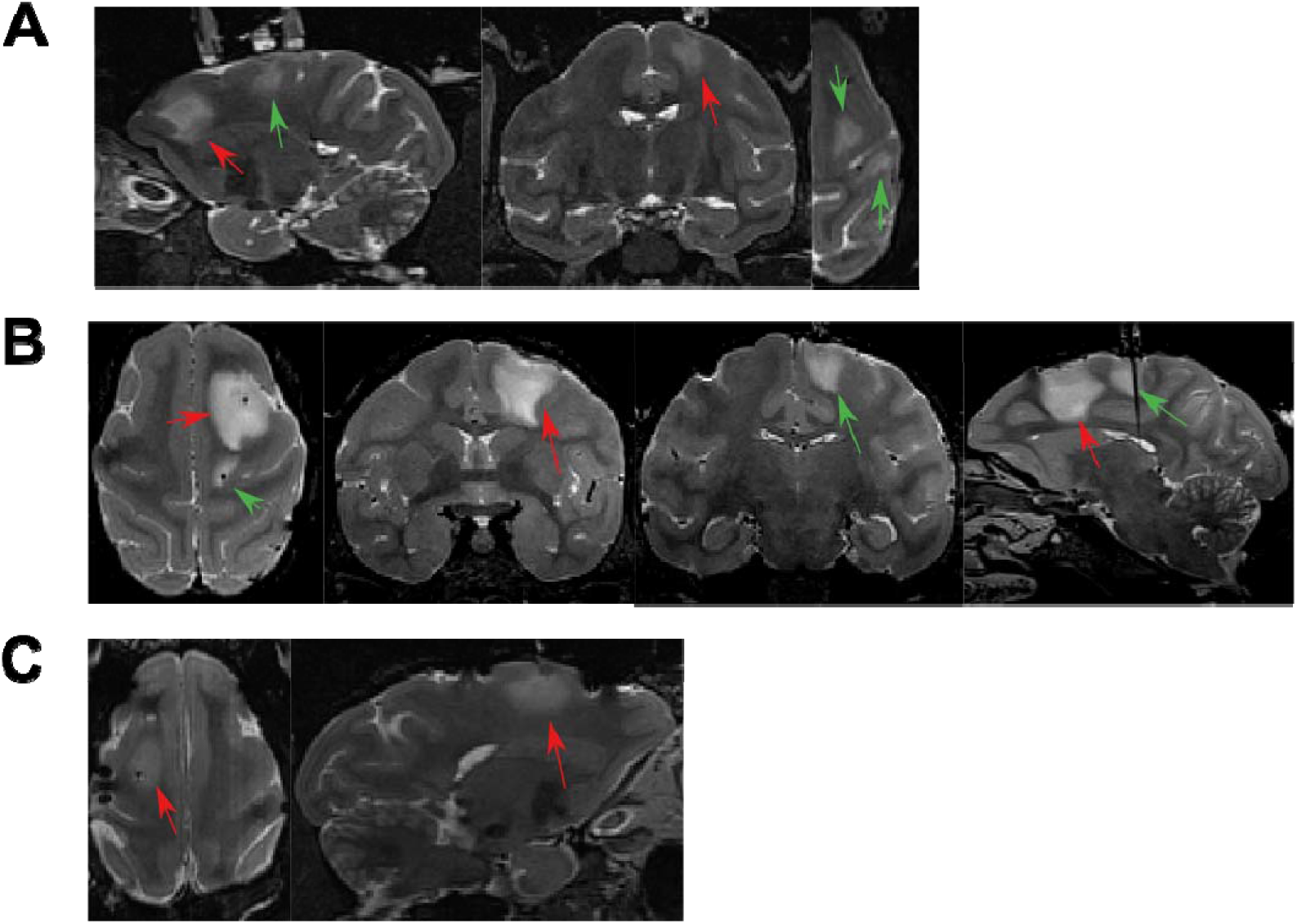
MRI findings in three additional cases. Similar T2 findings were found in 3 other cases. Case 2 (A) and 3 (B) had two regions underneath anterior (red arrows) and central craniotomies (green arrows), and case 9 (C) had one anomaly underneath the anterior craniotomy (red arrows. Note susceptibility-induced signal drops along the length of the electrodes in all scans as well as artifacts caused by titanium skull screws apparent in C.

Anomalous findings were rare following craniotomy surgeries conducted strictly with mechanical drills. There were six scans (across 5 animals) conducted within a week following these craniotomy surgeries, only one of which showed T2 hyperintensities (see Fig 2C). All others showed no anomalous findings despite electrode penetration at the time of the scans, indicating that the observed findings were unlikely to be caused by acute electrode placement for trajectory localizations (e.g., Fig 3A, 3B; cases 6 & 7).

**Figure 3:**
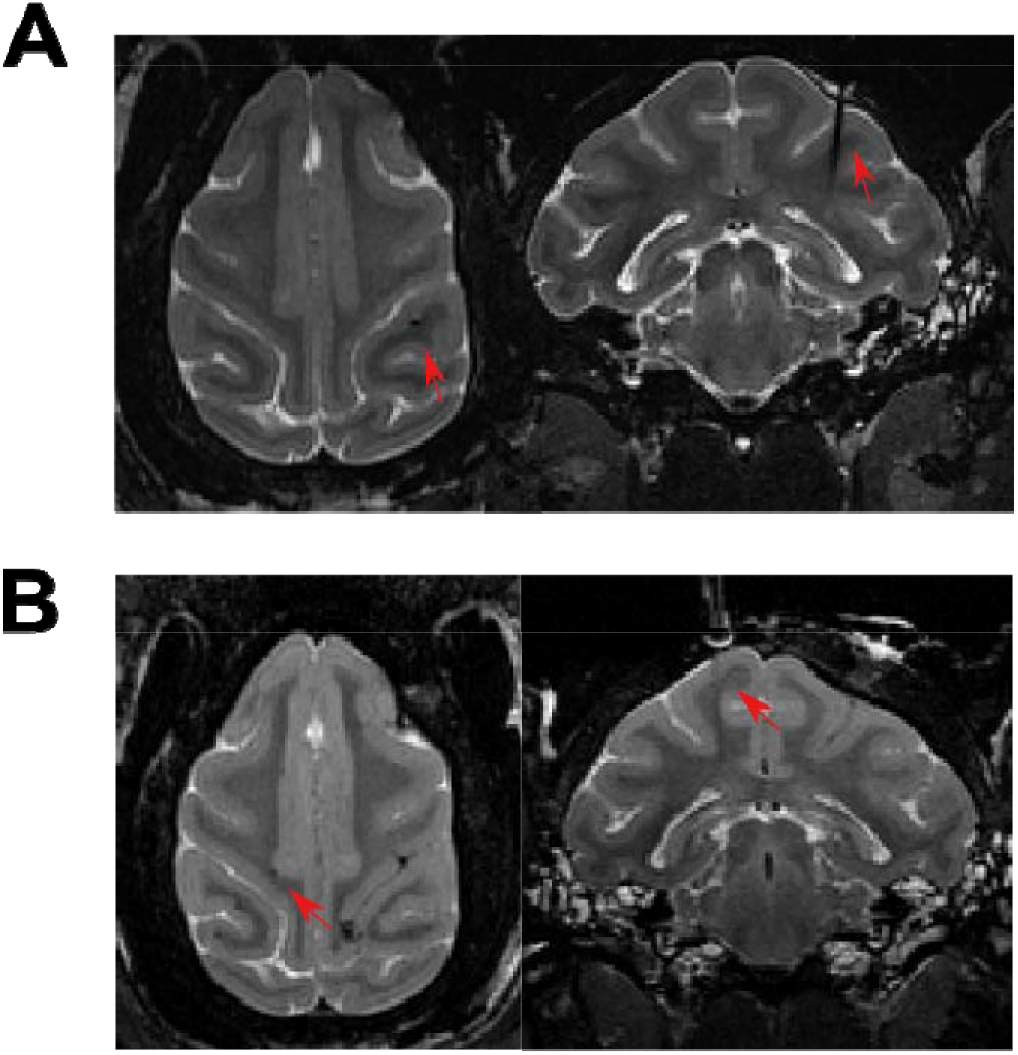
Negative MRI findings post-surgery. Exemplar negative findings underneath craniotomies (red arrows) in T2- weighted scans within 2 weeks from conducting craniotomy surgeries in cases 6 (A) & 7 (B).

In summary, six out of eight craniotomies done using piezoelectric drilling showed T2 hyperintensities (excluding the scan acquired more than two weeks following surgery), and five out of six done using electric drilling showed T2 hyperintensities (note the overlap in the use of the two tools in monkey P). In contrast, only one of thirteen craniotomies exclusively conducted with mechanical drills had underlying T2 hyperintensities.

## Discussion

Utilizing MRI-compatible implants in NHP research permits detailed whole-brain monitoring of animals following procedures and throughout the time during which the animal participates in research^12,17,18^. Our utilization of MRI-compatible implants enabled us to note a number of anomalies that may have otherwise gone undetected. T2 hyperintensities following craniotomies, particularly those performed with electric drills, were consistently found in our retrospective examination of craniotomy surgeries in eight animals. Inspecting T2-signatures in humans following craniotomy surgery, traumatic brain injury, and hemorrhages, hyperintensities found in our study are best compatible with a clinical interpretation of edema^2,14,19–24^. Our findings suggest that the mode of surgical drilling is responsible for the probability of edema occurrence at craniotomy sites. These anomalies resolve within a short period of time (e.g. two weeks in Case 1). Thus, there does not appear to be long-lasting structural (as seen in our T1/T2 scans) or behavioral effects. However, our findings raise the question of surgical tool recommendations in order to optimize neural tissue integrity in our research animals, which may affect experimental performance^24^, and especially as electrophysiological recordings may be conducted immediately post-surgery from tissue that may be physiologically compromised^25,26^.

Our results suggest that edema of neural tissue resolves within two weeks post-surgery, as indicated by a complete recovery observed in our structural scans. While we do not have histological data from our animals, data from the human and rodent literature suggest that resolution of neural tissue edema is of a similar timeline ^21,24,27,28^. One study showed that craniotomies using electric drills and trephines induced T2-hyperintensities and behavioral detriments to rats within two weeks post-surgery ^24^. Hyperintensities in animals with electric drill craniotomies were larger and were significantly reduced two weeks after surgery. However, in trephine surgeries, the affected areas were smaller and the resolution trajectories were heterogenous amongst subjects. The study suggested that these findings may be an effect of the disruption of nerve fibers and blood vessels integral to any craniotomy, in addition to trauma induced by vibrations or heat during surgery. Our contrasting findings in surgeries performed with mechanical and electric drills corroborate the timelines from this study, but suggest that trephines may induce effects on neural tissue dissimilar to the mechanical drills used in our study.

Post-operative MRI findings have been studied in humans (e.g. ^1,19^). Clinical examination of different tools and techniques is key for safer surgeries to reduce unexpected neural lesioning to the patient. While tools leveraged in animal work can be more diverse and less sophisticated than their counterparts utilized in humans, commonalities exist that may shed light on our findings here. It is difficult to identify the exact etiology of the edemas observed in our cases. Human studies have identified similar T2 hyperintensities following plunging incidents^1,3,19^. However, it is unlikely that our findings were due to plunging given the characteristics of the piezoelectric drill, the consistency of the findings, and the lack of any detectable deep migration of the tool (or dural tear) reported by the surgical team. Based on the findings in our scans, our data point to the possibility that use of either electric tool may be sufficient to cause an edema of the underlying tissue. Combining both tools in the same surgery did not indicate an additive effect; however, given our small sample size this conclusion is tentative. Conversely, surgeries that exclusively used mechanical drills were not likely to show post-surgical anomalies.

Contrasting signatures on T1 & T2 scans (Case 1) and recovery trajectory suggest that edemas underneath the craniotomies were induced either by mechanical forces imposed by an electric (piezo or otherwise) tool, or the heat generated by the tool. We suspect localized vibrations as a probable cause, because vibrations may reverberate through the skull and dura mater to affect neural tissue, a characteristic that resembles forms of mild traumatic brain injury^29^. Traumatic brain injury has been reported to cause reversible edema with similar T2/FLAIR profiles ^20–23^. Alternatively, thermal dissipation into brain tissue while drilling could also have caused the T2 anomaly; however, reports of lesions following thermal ablation by laser or ultrasound conveyed more focal, circular T2 findings that were sharply delineated from surrounding tissue^30–32^ and were long-lasting (observable more than two months following thermal application ^33^). It is also unlikely that thermal effects would be equivalent across the two types of electric drills as the piezoelectric drill includes an irrigation system that alleviates heat build-up during operation. Although less frequent, the occurrence of T2 hyperintensities in surgeries done solely with mechanical drills, which are unlikely to generate heat, also supports the notion that the anomalies were not due to thermal lesions. However, it should be noted that thermal effects reported in the literature were caused by mechanisms of heat application that are quite different from the ones associated with the tools used in our studies, which confounds a direct comparison with our T2 anomalies.

In summary, our results suggest that a more elaborate examination of surgical tools is warranted, especially given recent recommendations that embrace piezoelectric drilling as an ideal tool in macaque cranial surgeries to minimize risk for brain damage ^12^. This class of drill may prevent serious forms of plunging events with severe consequences to the animal, but may induce non-permanent lesioning despite an intact dural surface. While our findings suggest that there is no critical long-term clinical effect on brain tissue, it may be advisable to investigate pharmacological options for post-surgical edema attenuation^14,20,28^ or, more prudently, to delay neural recordings from cortical sites directly underneath craniotomies that were conducted using electric drilling for a few weeks. If this is not an option, mechanical devices appear to be a better choice to ensure the integrity of neural tissue underlying craniotomies.

## Acknowledgements

We thank the Princeton Laboratory Animal Resources staff for support.

